# Gradient Oriented Active Learning for Candidate Drug Design

**DOI:** 10.1101/2024.07.11.603160

**Authors:** Venkatesh Medabalimi

**Affiliations:** Independent Researcher

**Keywords:** generative active learning, directed evolution, black-box optimization, sample complexity, gradient descent, mRNA, minimum free energy

## Abstract

One of the primary challenges of drug design is that the complexity of Biology often comes to the fore only when proposed candidates are eventually tested in reality. This necessitates making the discovery process more efficient by making it *active*ly seek what it wants to know of reality. We propose Gradient Oriented Active Learning (**GOAL**), a technique for optimizing sequence design through active exploration of sequence space that interleaves performing experiments and learning models that propose experiments for the next iteration through gradient based descent in the sequence space. We demonstrate the promise of this method using the challenge of mRNA design as our guiding example. Using computational methods as a surrogate for experimental data, we provide evidence that for certain objectives, if one were restricted by the bandwidth or the number of experiments they can perform in parallel, increasing the number of iterations can still facilitate optimization using very few experiments in total. We show that availability of high-throughput experiments can considerably bring down the number of iterations required. We further investigate the intricacies of performing multi-objective optimization using GOAL.

## Introduction

One of the primary challenges of drug discovery and synthetic biology is that there are aspects to reality that are unknown and hence cannot be modelled. The eventual arbiter of truth is reality itself and many years of work towards discovery, the human effort and expenditure involved in the process can go completely in-vain due to the eventual failure of proposed candidates or methods in meeting the challenges and unforeseen obstacles that one gets to observe only at the end of the design pipeline (Dickson and Gagnon [2009], Rawlins [2004]). While this is always expected to remain a significant challenge given the complexity of Biology, with the advent of high throughput screening and emerging experimental techniques we have the opportunity to mitigate this problem to some extent by having a tighter and faster iterative loop in the discovery process that interleaves computational insight and experimental feedback from reality. It is thus a timely question to ask how can one bring these two parts of the process together so that there is the best possible information transfer between them while interleaving them so that the discovery process as a whole is expedited and benefits from doing so. We wish to have our computational insights drive our experimental curiosities and the results of these experiments should provide insights that further enable our computational models to ask better informed questions about reality. Such a feedback loop where *active learning* is baked into the process is the need of the hour.

Consider the example of mRNA therapeutic design, there are many aspects of an RNA sequence for which we do not yet have satisfactory realistic models. A sufficiently long RNA strand has many degrees of freedom due to potentially many interactions amongst its constituent bases that enable large variation in the structural configurations that can appear in an ensemble. A collection of RNA strands with identical nucleotide sequence can exist in an ensemble of possible secondary structures. Furthermore, for any secondary structure there are potentially multiple 3D conformations that it might exist in, which we do not yet completely understand let alone our ability to predict a distribution for this ensemble in given conditions(Vicens and Kieft [2022]). Besides their inherently high degrees of freedom, many aspects of reality can influence the properties and conformations of RNA in its medium, including downstream effects like its ability to produce desired target proteins, temperature, presence of other molecules like RNA-binding proteins in proximity, cell type-specific tRNA availability, translation factor abundances, mRNA decay factor abundances, ion concentrations and perhaps other things we don’t know about.

Given this uncertainty, it is entirely possible that reality might surprise us when our candidates are eventually tested. Besides, for usage at scale, there are often multiple criteria that we wish our candidate sequences satisfy at the same time. For example, unfailingly we want the candidate to be non toxic, thermally stable to facilitate storage, less vulnerable to degradation by hydrolysis, cleavage by endonucleases and exonucleases, have high degree of target protein expression and no off target effects. These demanding but necessary criteria make mRNA-design challenging(Metkar et al. [2024]) and make it necessary to make the most of interactions with reality through experiments we design along the way and optimize the discovery process to zero-in on promising candidates. We shall consider this application as a guiding example for showing a proof of concept of the technique we propose in this paper. At least, in hind sight, after the Covid-19 pandemic we consider this a biological problem ready for ML that is going to attract considerable in-silico ML research and hybrid ML-experimental biology research facilitated by high-throughput experimentation. Considering more elaborate approaches for sequence design provides an avenue for exploring the strengths and challenges of generative ML in biology, and an opportunity for experimental integration of generative ML while being grounded through interactions with reality.

We propose Gradient Oriented Active Learning (GOAL) as a promising technique for sequence design that zeros-in on desired candidate sequences through principled traversal of the enormous sequence space by iteratively learning function approximations for a measured property of the sequences. These models are then used to propose candidate sequences that are to be used for experiments in reality to get fresh grounded measures of the desired property. Crucially, the informed choice of which candidates to explore in each round is done using gradient flows at the currently available sequence candidates. We describe our method in finer detail in the Methodology section. We demonstrate that for certain desired properties this approach could be surprisingly effective in iteratively arriving at near optimal sequences in the search space. When facilities such as high-throughput screening are at our disposal this translates to minimizing the number of iterations in the computational modelling and experimentation loop. Even in scenarios when our experimental bandwidth i.e the throughput or the number of samples that can be experimented with at once is limited, we observe that considerable progress can be made in arriving at interesting parts of the search space by taking advantage of active learning over iterations. More broadly, it remains to be seen which kind of functional properties of sequences lend to such optimization by generative active learning and to what extent can expressive power and inductive biases of the computational models used have a bearing on the maneauvarability in the sequence space and effect the reachability to desired regions of the search space for those properties.

## Related Work

*Directed evolution*, recreating key features of evolution as a laboratory process to design interesting and useful molecules has long been proposed as a promising approach to over come the limitations of rational design Arnold [1998]. The key challenge in directed evolution is to make the entire process work in a much smaller amount of time while having an ability to produce interesting solutions with the desired properties. Generally, the search space of interest is huge and a randomly chosen candidate sequence or molecule is very likely devoid of properties that are sought and hence largely uninteresting, so the challenge is to direct, steer or nudge the evolution in such a way that the directed walk eventually reaches interesting parts of the space. A good starting point in the likely vicinity of desired target does help to begin with, but may not be necessary.

Romero and Arnold [2009] consider the problem of exploring protein fitness landscapes by directed evolution. In this approach one optimizes protein function over successive generations via random mutations followed by artificial selection or screening. They posit that this simple design algorithm can mitigate our lack of knowledge of how a sequence encodes for function and provides a reliable approach to engineering proteins with new and useful properties. More recently Fahlberg et al. [2023] proposes neural network extrapolation as a means to generalize to distant regions of the protein fitness landscape.

Embracing the idea of the entire scientific discovery process being data driven implicitly enables a richer hypothesis space that doesn’t leave out a whole lot on the table at the outset. One way to see what we seek to do is through the related lens of *Black-box optimization* which in itself is a rich discipline of study (Paria [2022].) A more recent line of attack towards black-box optimization is with the view of approaching it with the tools of generative-AI like diffusion models as proposed by Krishnamoorthy et al. [2023], Yuan et al. [2024] and Chen et al. [2024]. In a similar spirit Li et al. [2024] propose leveraging the power of diffusion models to capture data that has already been observed to proceed with optimization in a principled manner. Generative AI at large has shown significant potential in the design of bio-molecules like proteins, ligands and nucleic acids, an ability that holds immense promise for addressing pressing medical, industrial and environmental challenges. One of the first works to introduce generative optimization for DNA and RNA sequences using *activation maximization* (Simonyan et al. [2013], Yosinski et al. [2015], Mordvintsev et al. [2015]) and descent in the input sequence space of a learnt model was Killoran et al. [2017]. Castillo-Hair and Seelig [2021] propose Fast SeqProp a hybrid continuous/discrete model that uses position weight matrices to sample one-hot encodings corresponding to sequences in order to get improved performance. However, it is the synergistic integration of ML and experimentation(high-throughput or otherwise) that scales this effort as envisioned in Liu et al. that is the need of the hour.

Coming to the specific application of our interest, there are numerous considerations that should ideally go into the mRNA design challenge. A non-exhaustive list might include things as varied astranslation efficacy, avoidance of off target effects, avoiding toxicity, stability over a temperature range, customizing design so as to achieve potency in desired cell types and manufacturability Metkar et al. [2024]. In a step towards this Leppek et al. [2022] consider RNA-based therapeutic design as a combinatorial optimization of mRNA structure, stability, and translation. In a beautiful work, perhaps closest one to the application we consider, Zhang et al. [2023] proposes LinearDesign a principled mRNA design algorithm that concurrently optimizes both structural stability and codon usage. Addressing the problem of prediction leveraging a large collection of data, CodonBert by Li et al. [2023] is an LLM based model that has been trained using a comprehensive dataset comprising over 10 million mRNA sequences sourced from a broad spectrum of organisms, to predict properties such as protein expression and degradation.

An essential detour from taking a solely ML perspective is to think what can be done from a mathematical modelling and algorithmic perspective. If one were to fix a property like Minimum Free Energy(MFE) and consider the problem of minimizing it as a purely mathematical modelling endeavour, i.e constructing the coding sequence that codes for the given target protein but minimizes the consequent minimum free energy over all possible secondary structures of the sequence, the CDSfold algorithm by Terai et al. [2015] gives a polynomial time algorithm to find such a sequence. It finds a coding sequence(CDS) with the most stable secondary structure among all possible ones translating into the same protein. The dynamic programming based algorithm has time and space complexity of *O*(*L*^3^) and *O*(*L*^2^) respectively, where *L* is the length of the CDS to be designed.

Another important property of RNA to be aware of is its susceptibility to degradation through hydrolysis. This presents a problem in manufacturing and storage. Wayment-Steele et al. [2021] proposes to design RNAs to reduce mRNA hydrolysis by optimizing for double-stranded regions or minimizing unpaired nucleotides, which are not protected from in-line cleavage and enzymatic degradation. They observe that both sophisticated algorithms and rational design on Eterna (Lee et al. [2014]) provide constructs with low number of unpaired nucleotides(#UP) on average as opposed to conventional mRNA design methods. Elaborate structural data about RNA has only recently started to become available compared to other bio-molecules. Nevertheless, works like Wayment-Steele et al. [2022] have thrown light on the capabilities of commonly used RNA secondary structure packages and provided means to improve them by using high-throughput experiments. For the purpose of our work, in order to demonstrate a proof of concept of our paradigm we shall use the popular RNAfold package as a surrogate for providing us with what would ideally be high-throughput experimental data about properties like secondary structures of a sequence and the number of unpaired nucleotides in such secondary structures.

As a multitude of data starts becoming easier to obtain at scale - either due to new techniques (Dülk and Rouskin [2022], Cao et al. [2024]) or due to more public access for already existing data one should expect that ML methods that are fully data driven will be a competing if not preferable alternative for optimization to current ‘rational’ design approach. More so in the former case, with an ability to readily measure and assess newly generated samples in the input space for their properties while using a method like GOAL. Given the complex relationships between various mRNA properties such as unpaired nucleotides, stability, GC content, codon usage bias and mRNA structure, there remains significant potential for applying machine learning methods to develop generative models for mRNA design (Metkar et al. [2024]).

## Methodology

A typical loop in our procedure consists of three parts which are Annotate, Learn and Descend.

*Annotate* is the phase with real world experiments that takes a set of sequences and returns the experimental measurements of the desired property for those sequences. In the very first iteration we start with a set of uniformly randomly sampled sequences. For subsequent iterations the sequences to be annotated are supplied by the *Descend* phase from the previous iteration.

The *Learn*ing phase uses all the sets of sequences *S_i_* obtained and annotated over all the iterations *i* up until that point to learn a deep learning model *M_i_*which has a suitable inductive bias chosen for the property or function of the sequences we are trying to model.

*Descend* is the last phase of every iteration. In this phase we use the model learnt in the learning phase to descend in the input sequence space while keeping the learnt weights of the deep learning model fixed. We use each sequence from the most recent annotation as the initial seed for an individual descent, perform a gradient descent on the input sequence space in the near vicinity of the sequence to obtain a new candidate sequence to annotate. This gives us a new set of sequences to inform our understanding of reality when we annotate them in the next iteration. We shall call the number of sequences generated afresh by descent the ‘aperture’ of our method. The idea is that although the entire sequence space is humongous, perhaps its the case that for a well chosen inductive bias the deep learning model can produce a good function approximation that generalizes at least in the near-vicinity of the sequences that have been observed and annotated up until now so that when we descend from the most recent sequence set *S_i_*in this phase we obtain a better set of sequences on the fitness landscape (Wright [1932]). Crucially, the model *M_i_* has a path dependence that arises from learning its weights using all the collection of sequences until the current iteration and this influences the choice of sequences selected by the Descent phase to annotate.

(*Termination*) This process of annotate, learn and descend to produce new candidates is run for a pre-assigned number of iterations -or-up until the annotations for properties obtained from experiments fall in a desirable range for the requirements of our application -or- up until the annotations appear to saturate showing little progress over iterations.

It deserves a mention that for the functions we wish to approximate a *typical* uniformly randomly sampled sequence is likely quite boring and over-whelmingly unlikely to have a good desired value for the property of our interest. So when we sample a random collection of sequences more often than not they are all clustered around this typical poorly valued approximate level set, often with little variance in values (like the yellow set *S*_0_ in the illustration of GOAL in figure(2)). The challenge then for the ML model that learns a function approximation is to weave a sufficiently structured tapestry of the function landscape in the individual vicinity of these sampled sequences to be able to learn to descend with some confidence towards sequences that have better value for our property of interest.

**Figure 1:**
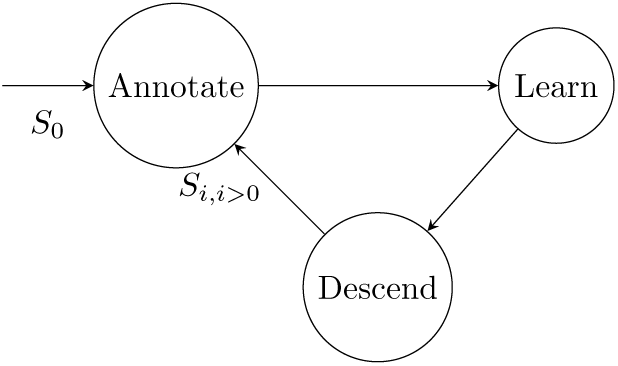
Phases of the optimization process, initial samples are randomly chosen while subsequent sequences for annotation come from the previous iteration after embeddings from the descent phase are rounded (*Rounding*) After the descent phase we have the embedding vectors for the new set of sequences *S_i_*_+1_ in the input space. To interpret them as a sequence of nucleotides(so as to annotate) we need to round each of the nucleotide embeddings to a one-hot encoding corresponding to some nucleotide. Throughout the process in each phase and across iterations we wish to keep our focus restricted to the nucleotide sequences that code for same amino acid at each position. To maintain this for each amino acid we map the embedding of the triplet of codons corresponding to it to the one-hot representation of the synonymous codon with the closest embedding. For the next iteration, these rounded vectors are added as the new sequences ready to be annotated (figure(1)). Note that unlike language or vision, biological sequences provide us the possibility of investigating through enquiring with real experiments(in vitro) the properties of freshly generated sequences that are produced in this manner-because they are physically meaningful constructs albeit even if they are unlikely to occur naturally. It is unlikely that similar attempts with language or vision would produce something coherent for humans to perceive let alone cognitively rate it.

**Figure 2:**
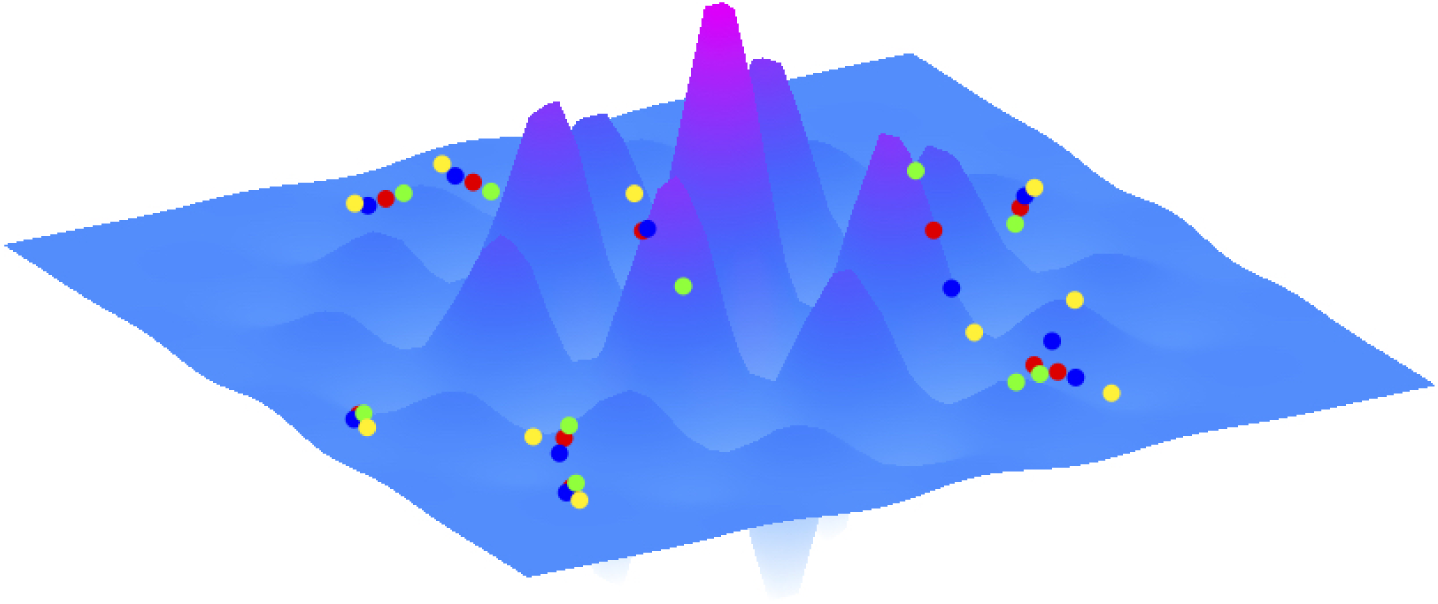
A simplified depiction of the progression of GOAL on a fitness landscape of sequences, sets of sequences generated in the process are depicted in different colors, *S*_0_*→S*_1_*→S*_2_*→S*_3_. Optimization here is illustrated as maximizing fitness. Successive models *M*_0_, *M*_1_, *M*_2_, *M*_3_ are learnt using the sequence sets *S*_0_, *S*_0_*∪S*_1_, *S*_0_*∪S*_1_*∪S*_2_, *S*_0_*∪S*_1_*∪S*_2_*∪S*_3_ respectively.

Let’s now focus on the specific problem of mRNA design and how each of these phases can be implemented. What properties of mRNA are to be considered and optimized for? How do we want the designed sequence to behaveOf particular interest in the context of drug design might be physical properties such as mRNA stability, its translation efficacy and perhaps most importantly in its interactions in vivo we want to avoid toxicity and off-target effects. Unfortunately, the author is constrained by lack of an ability to perform physical experiments to get real data. To mitigate this, we shall use widely used computational models as surrogates for what we need from real data in the Annotate phase. This will help us serve a proof of concept demonstrating the potential utility of GOAL in sequence design. It however has to be brought to the attention of the reader that there are a wide variety of computational tools available for RNA biologists (Fu et al. [2022], Reuter and Mathews [2010]) and the ability of these methods to closely approximate reality and their relative capabilities in doing so is a subject of study in itself. We shall use RNAfold from Lorenz et al. [2011], one such popular computational tool as our surrogate. As is the case with how the particular realistic context in which the mRNA is present can shape a physical experiment, the difference in computational surrogate used might effect the course of how the optimization and the demonstration of our technique would proceed.

We shall be primarily interested in minimum free energy estimates from RNAfold. In addition, we shall be using the number of unpaired bases in the MFE structure that RNAfold returns as a surrogate for the propensity of mRNA sequence to decay. Also, one other criteria that has been purported to correlate with translation efficacy is the Codon Adaptation Index of the mRNA. When we need it, we shall use the relative codon usage rates observed in the human deltoid tissue obtained from https://dnahive.fda.gov/ as our reference when we consider a higher CAI as desirable. We investigate to what extent our method GOAL can help us achieve desired properties and what happens when we optimize them in tandem.

For the Learning phase we observed that an architecture akin to Residual Convnets (He et al. [2016]) served the purpose of being able to efficiently learn function approximations. However, it is very much possible that there are architectures with much better inductive biases corresponding to each of the desired properties of mRNA that can help model the function better and extrapolate further from the vicinity of the current set of sequences *S_i_*. We elaborate more about the details of our experiments and the accompanying results that address the questions we seek to explore in the next section.

## Experiments and Results

In the experiments we set up, we wish to investigate the following - Can the sample complexity of optimization by learning be decreased significantly using GOAL in comparison to a static approach that uses randomly sampled data and seeks to optimize using this data without going back to actively procure more data? Within the framework of GOAL, we investigate if using a large aperture as enabled by high-throughput data expedites the optimization in comparison to a small aperture? Furthermore, we are curious how the number of iterations to converge and the quality of the solution arrived at is effected by using a small aperture i.e a smaller number of samples per iteration.

It is expected that the outcome of this investigation would rely on the nature of the property or objective(s) we are trying to optimize. For our proof of concept, we shall primarily consider Minimum Free Energy(MFE) as the objective to minimize using GOAL. Later in this section, we investigate what can be done when we seek to optimize multiple objectives using GOAL. Other objectives of interest can be as diverse as-number of unpaired bases in the secondary structures, Codon Adaptation Index of the sequence, some well chosen measure of toxicity or a measure of likelihood of mis-translation. One can also place hard constraints that seek to avoid certain sequences, for example those that have a high likelihood of secondary structures (Kikin et al. [2006]) with g-quadraplexes potentially causing mis-translations due to frameshifting (Yu et al. [2014]). Likewise, as is the requirement one might want to avoid sequences which produce secondary structures that have very long stretches of dsRNA that are known to trigger global shut down of cellular mRNA translation after sensing by protein kinase R (Hur [2019]).

Now we give further details about our experimental set up. The initial set of sequences to annotate, *S*_0_ are generated by sampling sequences uniformly randomly from the set of all sequences formed by using any of the synonymous codons for each amino acid in the sequence coding for the target protein. These are then annotated for their MFE using RNAfold (Lorenz et al. [2011]). All the subsequent sequences the algorithm ever requires to annotate are obtained as a result of the descent phase in the previous iteration.

When considering a computational surrogate under limitations on resources at one’s disposal a natural trade-off one runs into is between the time it takes to compute the MFE structure and the realistic nature of the underlying model for MFE computation. One can look back to traditional score-based dynamic programming algorithms for a simple reference. The underlying DP based MFE calculation is simple and we can compute minimum free energy estimates for a given RNA sequence using Nussinov’s algorithm in *O*(*n*^3^) time where *n* is the length of the sequence (Nussinov and Jacobson [1980]). However, this doesn’t scale well if we were to use it for generating large datasets comprising of sequences of a long length *n* for example, like the spike protein of SARS-Cov2 which is 3819 nucleotides long. Nevertheless, we can observe that the structure of underlying DP is parallelizable and we can avail a GPU to bring the effective complexity for our practical purposes down to *O*(*n*^2^). This produces a scalable way to generate MFE data to bootstrap. But, one might legitimately object that the above computation is too simple and the underlying computational model doesn’t attempt to capture reality in a satisfactory manner. For example, it lacks a detailed thermodynamic model and empirical parameters that programs like RNAfold incorporate on top of considering the stability conferred by base stacking, and allowing for possibilities like wobble pairs. Alternatively, we can use a readily available popular folding algorithm like RNAfold as a surrogate for reality. Given the inherent time consuming nature of these folding algorithms whose readily available implementations don’t run on GPU, we avail multi-core machines to harness data parallelism while performing our annotations. This provides us with an order of magnitude speed up that compensates for the time consuming nature of each individual computation on the long sequence.

The architecture of the deep learning Model that we use to learn a function approximation of MFE is shown in figure(3). We use what might be called a version of Residual Conv-net arrived at after much tinkering to understand the manner in which the choice of the model effects the entire process of learning and subsequent optimization using the model. It is worth mentioning that it is by no means a settled question that this is the best network architecture one can consider, nevertheless it suffices to demonstrate a proof of concept of our technique. Input sequences are represented using one-hot encoding where each base is coded by a 4 *×* 1 one-hot vector.

**Figure 3:**
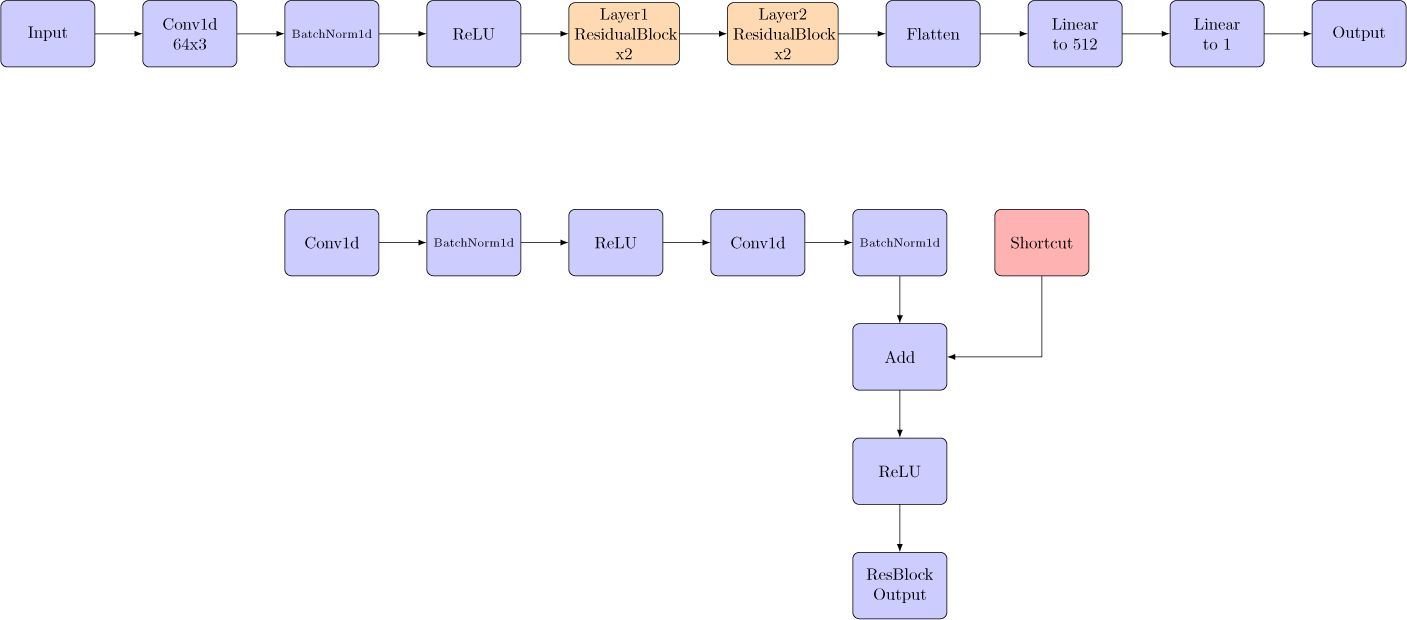
The architecture of our ResNet model with a detail of the Residual Block.

Learning rate is set to 0.001 in the learning phase while the network weights change. In the descent phase when the change happens in the input space it is set independently to a suitable value as per the objective under consideration. All the models are learnt using MSE loss but we found the use of SoftRankLoss equally effective.

During descent in the input space, for each triplet *x_i_*_:*i*+3_ in the nucleotide sequence, the 4 *×* 3 input vector corresponding to each codon is projected to the closest codon that codes for the same amino acid. The distance of *x_i_*_:*i*+3_ to the 4 *×* 3 one-hot encoding of candidate triplets corresponding to different synonymous codons is measured by the Frobenius norm, 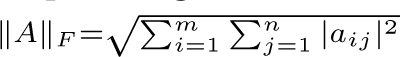 So *x_i_*_:*i*+3_ projects as follows

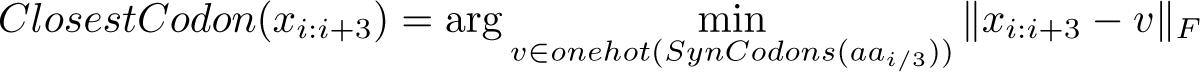

where *aa_i/_*_3_ is the required amino acid at the *i/*3*^th^*position of the sequence for the target protein.

We take the task of designing mRNA for SarsCov2 spike protein with the lowest possible minimum free energy when the MFE for a sequence is computed using RNAfold. The optimal sequence as given by CDSfold algorithm (Terai et al. [2015]) provides a good alternative to contrast the solutions we arrive at using GOAL. We observe that we are able to arrive at sequences that are within 89 percent of the minimum free energy possible as is known from CDSfold algorithm. We observed that with an aperture of 25k samples i.e what one might consider high-throughput annotation, the method converges to the local minima that is 89 percent of the global optimal within as few as 7 iterations.

On the other hand, we observed that one can arrive at considerably low MFE sequences even while using a small aperture but running the procedure for a longer number of iterations. Somewhat surprisingly, even with a very low aperture at only 50 annotated samples per round the method manages to get to roughly 80 percent of the global optimal MFE but does so in about 30 rounds. This comes to a total of merely 1500 sequence annotations, showing us the power of exploring and optimizing in the sequence space *actively* in an informed manner. This is demonstrated in figures (4 & 5). The sample complexity of annotation, the proxy for cost is the number of physical experiments one would have to run during our learning procedure and is given by the number of iterations times the aperture. So there’s a visible tradeoff between the size of aperture of our active learning trail through the sequence space and the number of rounds it takes to converge, consequently there might be a sweet-spot in terms of choosing an aperture large enough so as to expedite the time it takes for optimization and yet favorably small enough so as to minimize the cost incurred in the process without effecting the quality of the solution.

**Figure 4:**
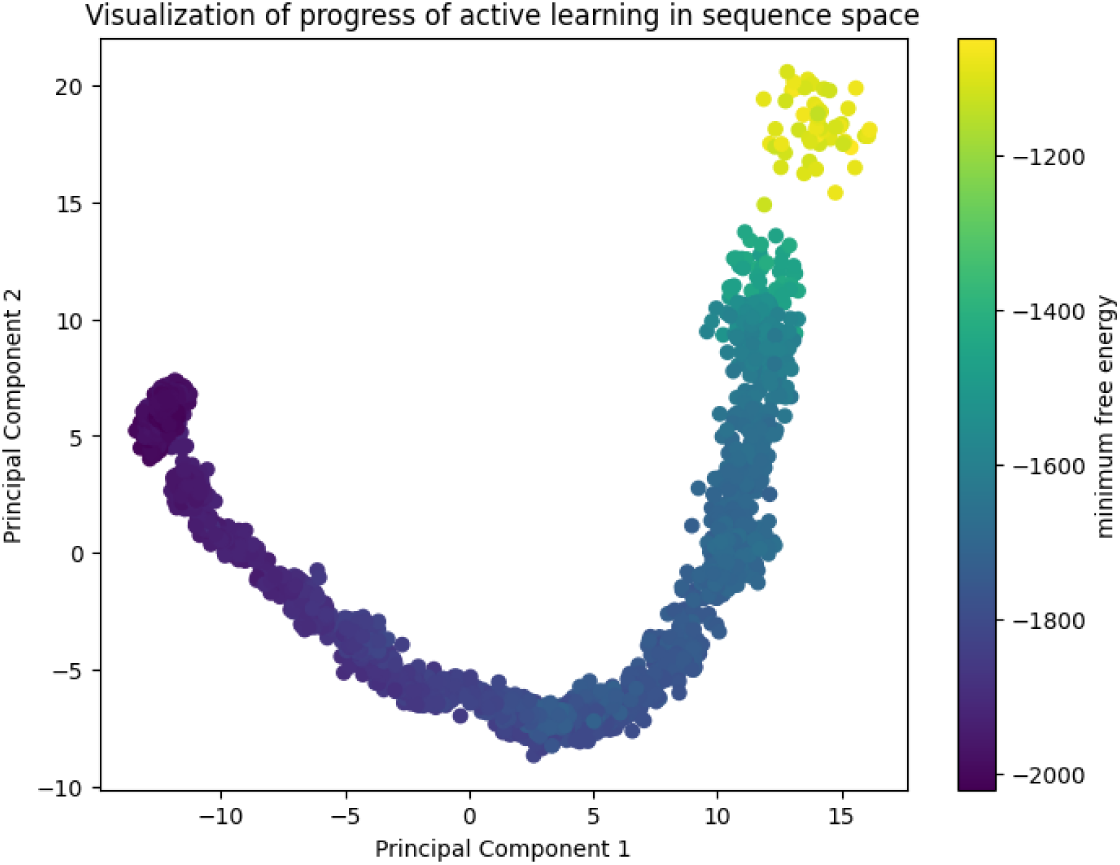
depicts the progression of GOAL as the population of sequences pierces through the sequence space to eventually arrive at a collection of candidate RNA sequences with a low minimum free energy. The depiction is in terms of the top two principal components of the set of sequences that constitute the entire optimization trail.

**Figure 5:**
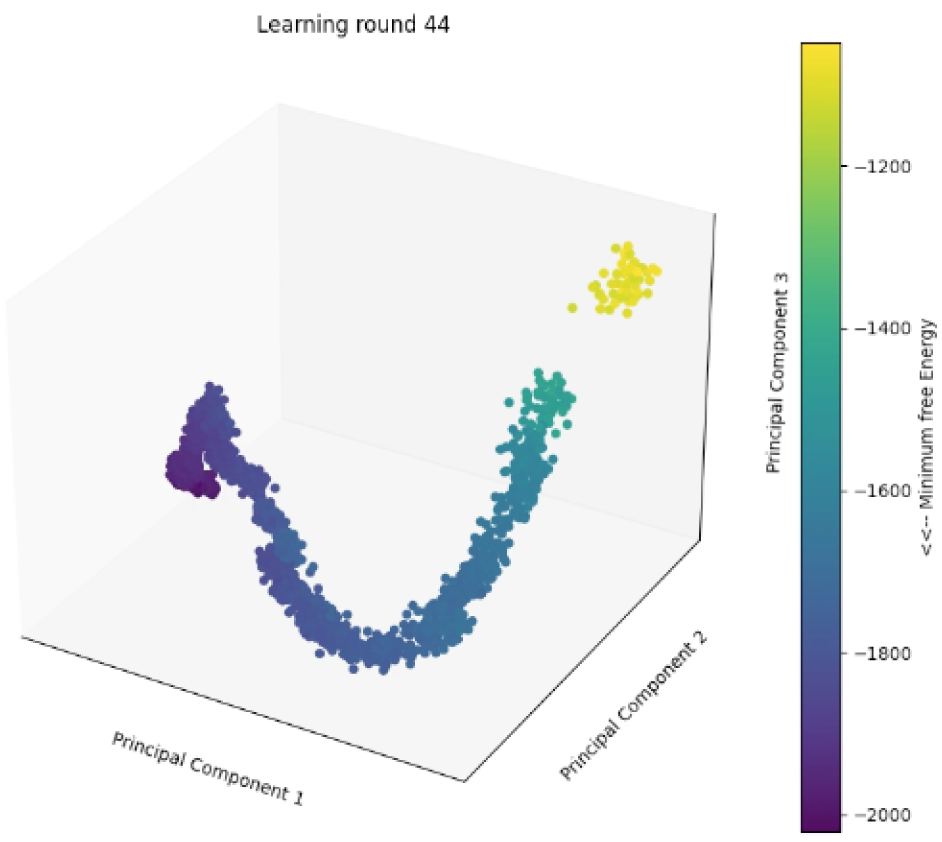
depicts the progression of GOAL as the population of sequences pierces through the sequence space to eventually arrive at a collection of candidate RNA sequences with a low minimum free energy. The depiction is in terms of the top three principal components of the set of sequences that constitute the entire optimization trail.

One can observe that during this learning procedure the population of sequences become less diverse and get closer to each other over time. Figure(6) depicts the average pairwise distance between the collection of sequences freshly obtained at the end of each round. Figure(7) depicts the average inter sequence distance between the seeding sequence and the sequence generated upon descent from it using the learnt model. Figure(11) shows the MFE structure of one of the RNA sequences from the final collection arrived at as a result of optimization and for contrast, figure(12) shows the global optimal under the same energy model arrived at using the CDSfold algorithm.

**Figure 6:**
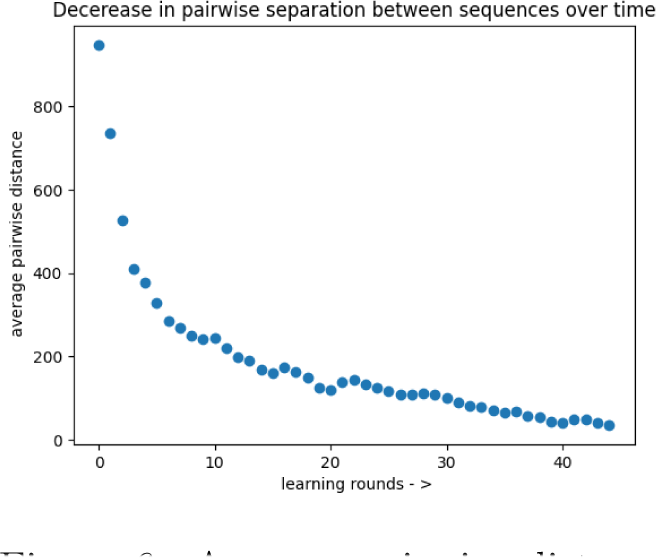
Average pairwise distance between sequences in the population over the iterations

**Figure 7:**
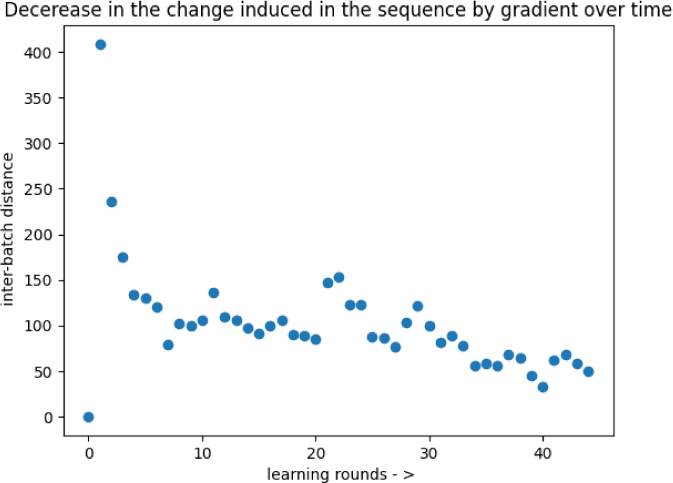
Average inter sequence descent distance between corresponding pairs of sequences over iterations

**Figure 8:**
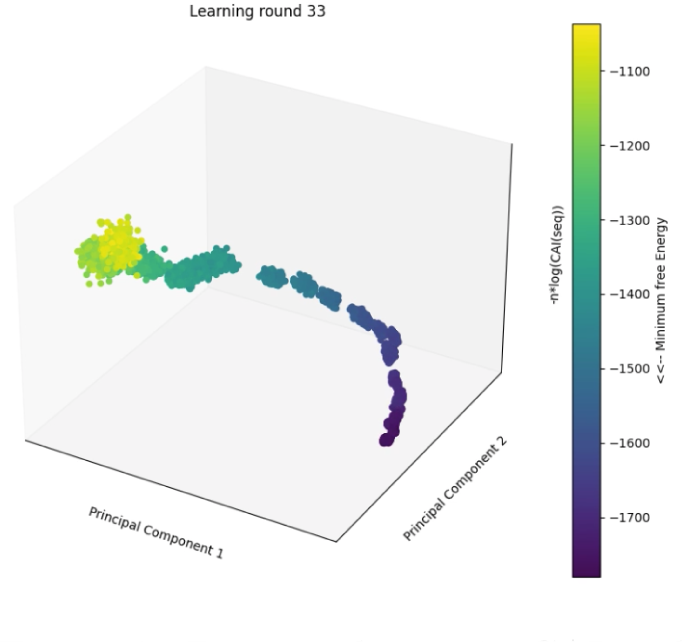
Depicts change in CAI and MFE during progress of optimization of 3 part objective comprising: CAI, unpaired bases in MFE structure and MFE weighted equally.

**Figure 9:**
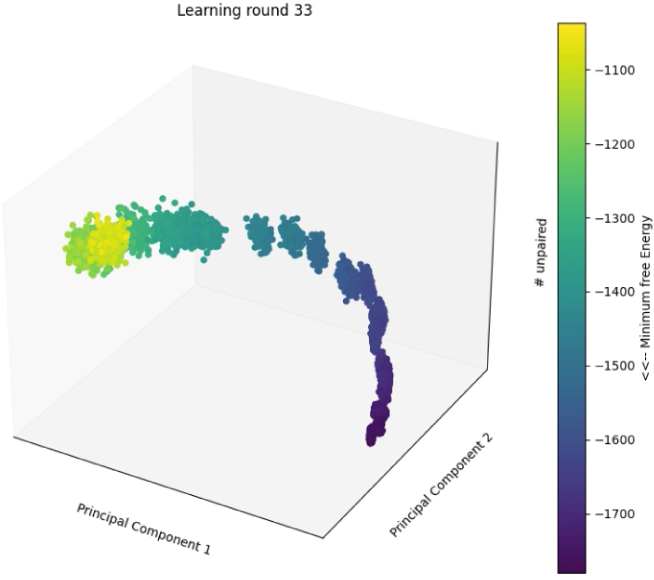
Depicts change in unpaired bases and MFE during progress of optimization of 3 part objective comprising: CAI, unpaired bases in MFE structure and MFE weighted equally.

One of the primary challenges in the development of mRNA therapeutics has been to understand and regulate the characteristics of mRNA that trigger innate immunity. One such characteristic is presence of long double stranded RNA helices, where it is known that dsRNA longer than 33 base pairs can trigger global shut down of cellular mRNA translation after sensing by protein kinase R (Hur [2019]). Intriguingly, we observe that the longest double stranded stretch in the optimal structure arrived at using GOAL(depicted in figure(11)) is 17 bases long while for CDS optimal structure(in figure(12)) it is 28 bases long. We speculate that this might be attributable to the nature of inductive bias that Residual-ConvNets bring in. Wayment-Steele et al. [2021] observe that the sequences arrived at using their RiboTree algorithm and LinearDesign from Zhang et al. [2023] also don’t have long stretches of double stranded RNA helices. However, there are potentially many more aspects to understanding how sensing mechanisms associated with innate immunity recognize RNA structure and trigger response mechanisms. Needless to say, these aspects therefore need to be under careful consideration for any design endeavour (Kato et al. [2006], Patel et al. [2012], Peisley et al. [2011]). The author is ignorant of these important aspects of reality that need expertise in immunology and wishes to bring this to the attention of the reader.

### 4.1 Multiple Objectives

Until now we have focused on a single objective, having multiple objectives to consider and attempting to optimize them simultaneously can potentially present us with many challenges. There will be choices to be made and decisions to be taken along the way that shall be primarily determined by our requirements in terms of those objectives and the extent to which these individual objectives align with each other. We are curious how one can extend a framework like Gradient Oriented Active Learning to multiple objectives. Some of the choices one faces along the way include

- Can we transform the problem into a single objective problem by weighting the individual objectives in some good proportion according to the relative priorities we place on them? Figure(10) depicts the optimization trail for such a joint objective that seeks to minimize an equally weighted objective that comprises of the number of unpaired bases in the MFE structure and the MFE. Figures (8) & (9) depict progress during optimization of a 3-part equally weighted objective comprising Codon Adaptation Index, number of unpaired bases in the MFE structure and the MFE, where the contribution from CAI is taken as *n* log(*CAI*(*s*)) to make it commensurate with other parts of the objective just the way Zhang et al. [2023] do. Alternatively, an arguably more judicious choice of these relative weights given to parts in the objective might be made from how these objectives interact in reality. For example, while considering MFE and number of unpaired bases in the MFE, a more elaborate way to model reality through the objective might be arrived at by considering probability densities of secondary structures(determined by free energies) in the en-semble approximated by Boltzmann distribution and corresponding decay rates determined by number of unpaired bases.
- We can choose to minimize multiple objectives simultaneously by computing an appropriately aligned displacement vector from the gradient updates suggested by individual optimizers as is proposed for example by Sener and Koltun [2018]. We find that in this case the optimization might be heavily influenced by a single weak and(or) discordant model. This could be arising from a small gradient vector that is also potentially aligned in a different direction caused by the nature of a single objective at the samples currently at our disposal.
- Alternatively, one can consider optimization by interleaving the individual optimizers as if it were children taking turns to play with a string of beads in their own way. In this case however, we observe the process is highly swayed or influenced by the optimizer that has comparably larger gradient and consequently takes the longer strides for the current population. Thus when interleaving the models, its not immediately clear what is a principled way to choose the relative step sizes for each optimizer, for it would need an understanding of the fitness landscapes of these objectives. This also makes a case for exploring other novel ways to steer or direct the evolution of population of sequences available.

**Figure 10:**
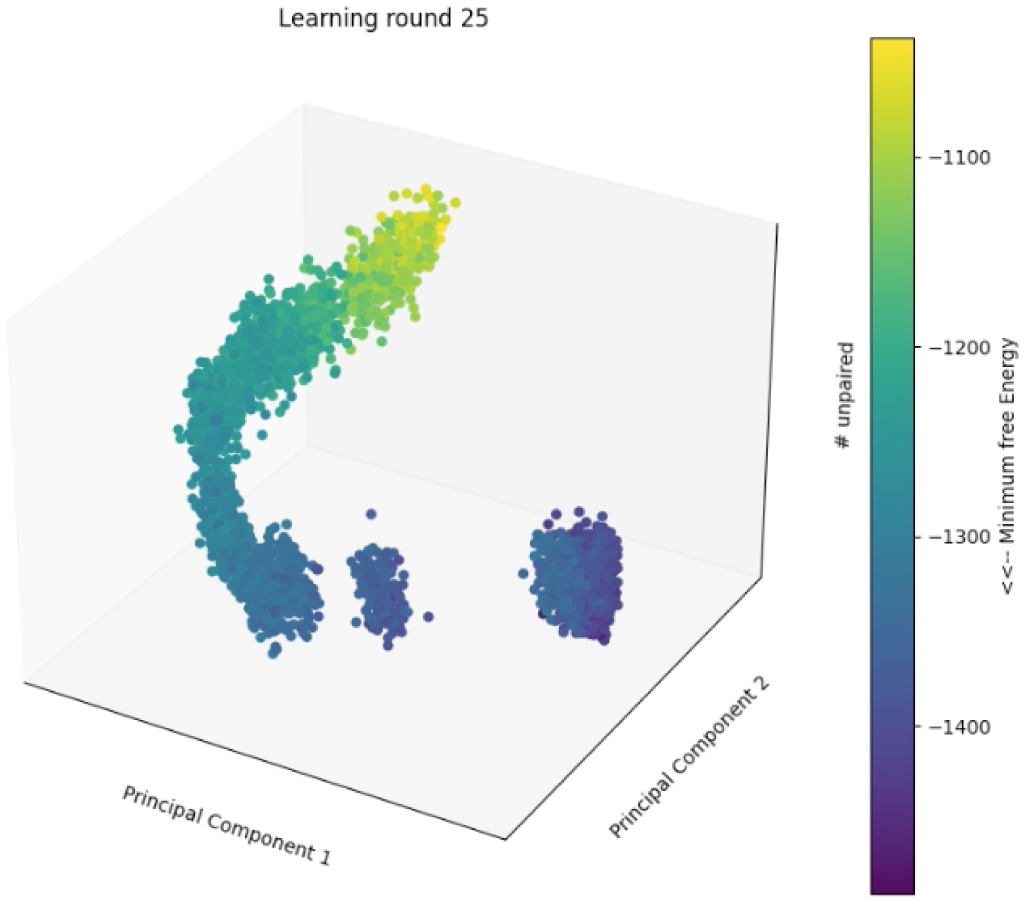
depicts the progression of GOAL during joint optimization of number of unpaired bases in the MFE secondary structure and MFE itself when guided using a single objective function that gives equal weightage *α* = 0.5 to them, number of unpaired bases goes down along z-axis and the color code gives MFE. The depiction is in terms of the top two principal components of the set of sequences that constitute the entire optimization trail.

**Figure 11:**
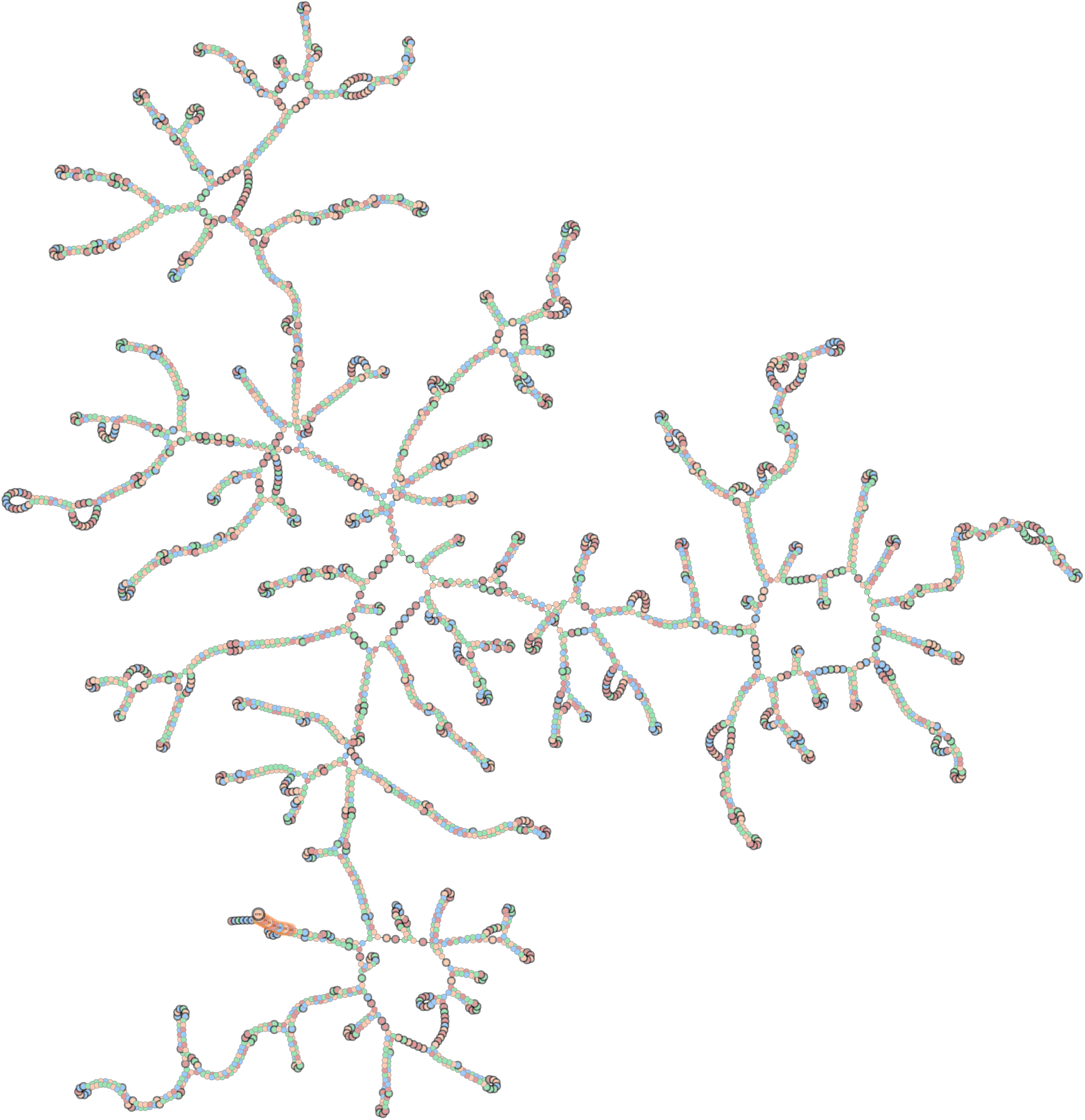
A force directed plot of the MFE structure of the best sequence in the eventual sequence set S obtained by GOAL for the SARS-Cov2 spike protein rendered using graph-tool (Peixoto [2014])

**Figure 12:**
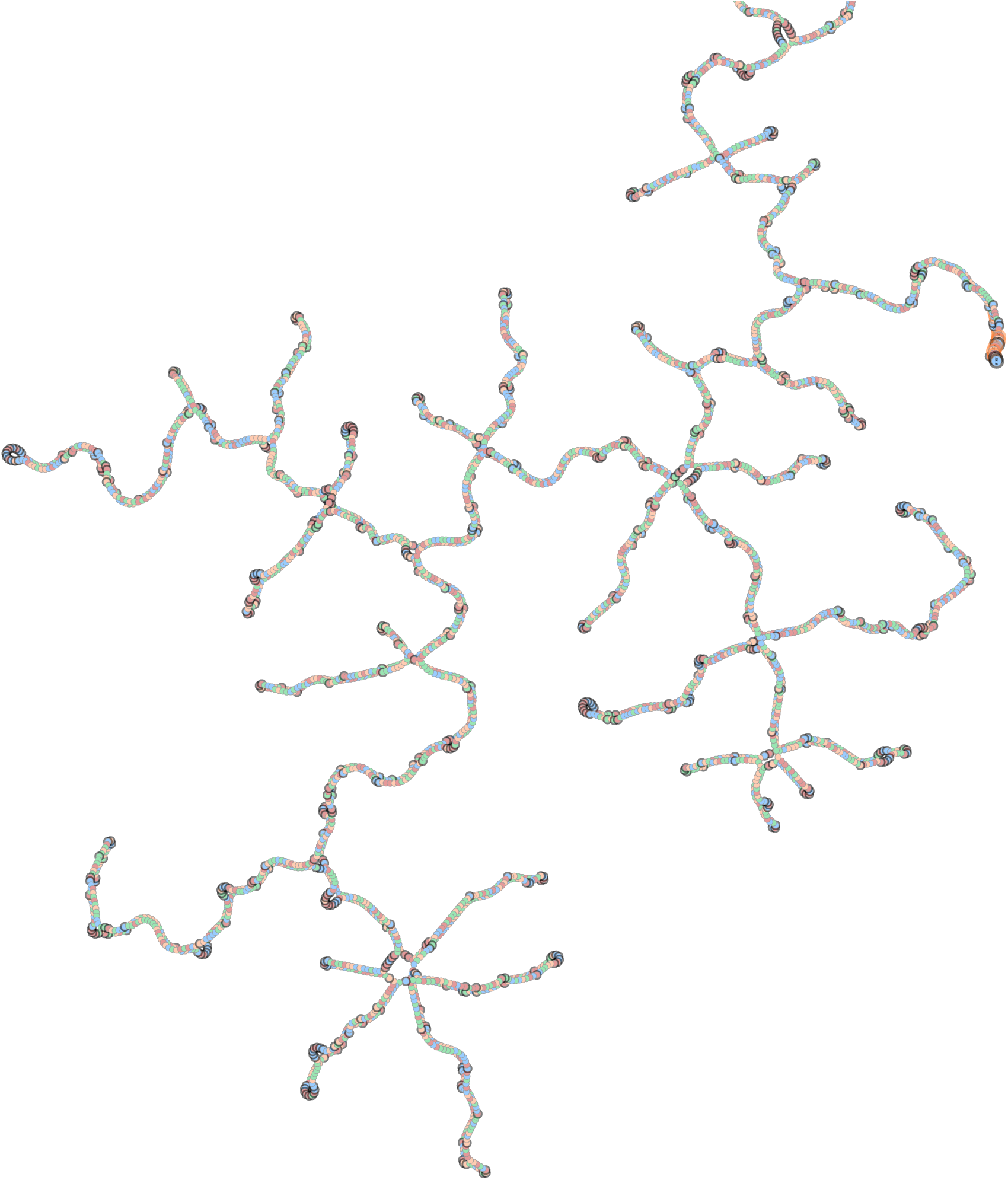
MFE structure of the optimal sequence given by CDSFold algorithm

Note that these choices and observations are merely a sample to give a glimpse of the complexities involved with multiple objectives and not meant to be exhaustive by any means. Consider for example the multi-objective task of minimizing MFE and number of unpaired bases in the MFE structure, if we use the same residual ConvNets from figure(3) to learn individual models for these objectives and use GOAL on their input space, the learnt models suggest the following dissonant strategies. Minimizing MFE causes an increase in GC content while minimizing #UP causes an increase in UG content. This happens presumably because when starting from a pool of uniformly randomly sampled sequences, the MFE minimization wishes to benefit from allowing for more GC base pairs that are relatively more stabilizing. While for minimizing #UP, starting from a pool of uniformly randomly sampled sequences, increasing UG content can provide the optionality of facilitating both canonical and wobble pairing to decrease the number of unpaired bases. This dissonance of misaligned strategies can lead to inefficient local optima even in comparison to using a single optimizer. Perhaps surprisingly, we observe that the solo MFE minimizer eventually manages to arrive at sequences with definitively lower number of unpaired bases in their MFE structure than the #UP minimizer does by itself. More generally, as is the case with first-order methods a problem that one might anticipate is that even when the descent direction chosen might not be ideal, due to the path dependence inbuilt in GOAL depending on how the landscape is shaped the process might make the population double down on the path to associated local optima as iterations proceed. As an example, as #UP optimizer chooses to increase UG content the chances of it heading into high GC content space (where we know better solutions do exist as shown by the MFE optimizer) go down due to its new found evidence upon descent reinforcing or affirming its implicit hypothesis that #UP can be decreased in a certain manner.

## Discussion

As observed earlier, single objective optimization using GOAL might take you to novel parts of the sequence space producing interesting local optima. GOAL might be able to do so using relatively few number of learning iterations or a small number of physical experiments by actively learning a model and proposing experiments in reality that are informed by this learning. While it is indeed the case that GOAL can produce such interesting solutions there is no guarantee that this is always the case and the path dependence inherent in learning can create a propensity to reinforce descent towards suboptimal solutions. In addition, the primary constraint against actively learning from local information is perhaps the complexity present in the nature of a property’s variation on the sequence space. Much like it is an impedance for evolution itself (Kaznatcheev [2019]), Complexity is perhaps the ultimate constraint against optimization using such evolution like techniques.

It is an interesting problem to investigate principled methods to better steer optimization using GOAL. More so, in the context of multiple objectives the risk of falling prey to simplistic or naive coherence in the presence of misalignment in the objectives is present. This naive coherence can lead the process to arrive at shallow optima or in other words- low complexity solutions. Indeed it might be the case that being guided or nudged by a thoughtful designer might be a useful way to mitigate the problem of falling prey to the solution that might be arrived at from the limitations of using only local information on the fitness landscape(s). Nevertheless, who is to say what is hidden in as yet unexplored parts of the sequence space for complex properties that are hard to comprehend and tease apart? One might even argue Evolution is merely trying to find a way and not to optimize and yet - has brought to bear all the rich variety around us!

## Notes

### Competing Interest Statement

The authors have declared no competing interest.

